# Decontextualizing Big Data for a Better Perception of Macroecology

**DOI:** 10.1101/059915

**Authors:** Christian Mulder, Giorgi Mancinelli

**Affiliations:** National Institute for Public Health and the Environment, Bilthoven, The Netherlands; ECOTEKNE, University of Salento, Lecce, Italy

## Abstract

Fish species are charismatic subjects widely used for ecological assessment and modelling. We investigated the influence of electrofishing in an attempt to illuminate the extent to which datasets might be merged together. Three American Midwestern regions in Ohio were chosen as study area and the changes in the size-biomass spectra of more than 2000 fish assemblages were analysed. These communities behaved differently according to the sampling method in conjunction to the morphology of the investigated streams and rivers, as shown by decreasing predatory species and lowering of allometric slopes. There are here several lines of evidence indicating that the chosen sampling method, as determined by different habitats, acts as a pitfall and strongly influences the allometry of fish spectra. In the ongoing data-rich era, our results highlight that macroecological investigations, often performed by machine-learning systems without considering the different procedures adopted to collect original data, can easily produce artefactual allometric scalings.

## Introduction

Big Data and eScience strongly facilitate data-intensive research across disciplines (Hey et al., 2009), with several domains ranging from genetics (www.ncbi.nlm.nih.gov/guide/data-software) and socio-economic sciences (www.icpsr.umich.edu/icpsrweb/IDRC), through services (www.naturalcapitalproject.org) and environmental information (e.g., mrdata.usgs.gov) up to macroecology (www.ngdc.noaa.gov/ecosys/ged.shtml), invasive species (www.issg.org/database), and biodiversity (www.gbif.org), with hundreds of other websites-often not freely accessible-like those for charismatic taxocenes (www.fishbase.org and ebird.org), vascular plant traits (www.try-db.org), and food webs (globalwebdb.com).

This wealth of Open Data and the improvement of machine-learning packages enable large-scale investigation (Sibly et al., 2012; Hampton et al., 2013). However, it is not solely the identification and verification of macroecological laws which are challenging in a data-rich era, but also the compilation from publicly-accessible portals of new databases with information collected by strictly standardized sampling protocols.

Recently, Raffaelli et al. (2014) correctly posed the question “*How might one design an ecosystem-level research programme that utilises Big Data approaches and frameworks?*”. Their question is far from rhetoric. In many investigations, most care is devoted to taxonomic pitfalls; however, when merging records with different protocols together, greatest care is provided to environmental parameters (temperature, pH extraction, molar vs. atomic ratios etc.) but in contrast to vegetation surveys, far less attention is given to data compilation when records of faunal populations assessed with different sampling methods are merged together.

Given the widely discussed universality of allometric scaling, Big Data can allow the detection of such sampling pitfalls. Size-biomass spectra enable us to plot the body size-binned biomass distribution of single species in a food web, and to model the resulting allometric scaling according with the sampling procedure. The interpretation of biomass spectra is intuitive (Mulder & Elser, 2009): the linear regression slopes fitted across the spectra indicate if biomass changes with increased body size, with steeper slopes indicating a negative response of the smaller-sized organisms and a positive response of the large-sized organisms, whilst shallower slopes indicate the opposite. This conceptual model is expected to be a general statistical tool: if scaling is universal, allometry must be unrelated to any sampling method.

We will investigate possible effects of different electrofishing sampling methods on the allometric scaling of freshwater fish assemblages by integrating theory and empirical approaches. Assemblages have been quantified in terms of fish population sizes and binned biomass in combination with information regarding diet and life-history attributes obtained from FishBase by comparing the size-biomass spectra between sampling methods. By verifying the connection between different electrofishing procedures and allometry, we will analyse if (and if so to which extent) a sampling methodology might affect allometric scaling. Analyses will be organised along the following chain of relationships:

1. The local hydrogeomorphology of a water body forces the choice of specific electrofishing procedures;
2. The electrofishing method allows to define the composition of a fish assemblage;
3. Body-size determines population size (hence, biomass) within a fish assemblage;
4. Population sizes and biomass determine material flow rates across trophic levels;
5. Material flow rates determine size-biomass spectra and hence allometric scaling.

## Study Area

The most widely used procedure to obtain robust fish abundance and distribution data is pulsed direct current (DC) electrofishing (Simpson & Reynolds, 1977; Flotemersch, 2001). Several sites across Ohio, USA, were sampled between 2000 and 2007 (www.epa.gov); we used 41,070 records of fish counts and body-mass estimates to build a prominent dataset on 134 freshwater species and 36 hybrids. The resulting dataset comprises 2051 locations (Figure 1 upper left): 406 sites are rivers sampled by boat-mounted electrofishing (straight electrode array: Method A), 794 sites are larger streams sampled by wading electrofishing (tote barge: Method D), and 851 sites are smaller streams sampled by longline electrofishing (longline: Method E).

The structures of fish assemblages were assessed using fresh M (species body-mass averages), N (counted fishes), and B (species-specific biomass: log(B) = log(N) + log(M)) data. This procedure produced log(N), log(M), and log(B) attached to each node. Most nodes are trophically highly connected (Figure 1), suggesting cascading effects as soon resource nodes will disappear. After lumping fish species with comparable site-specific average weight together, the total biomass for all taxa occurring at a given size bin was plotted at the middle of that size bin.

To compute size-biomass spectrum slopes using bins of constant linear width, the range from the smallest log(M) to the largest log(M) in all assemblages was divided into 14 equal size bins and for each bin the log of total biomass was computed. All values were regressed against log(M) at the centre of each bin (zero observations were excluded) and the log(B) = a⨯log(M) + b was fitted to each site (confidence interval 95%) and along the binned log(M). Allometric results were compared with sampling methods by means of nonparametric analyses (Mantel & Valand, 1970).

## Results and Discussion

In Figure 2 the regression slopes were discretized and assigned to 14 bins determined by the Freedman-Diaconis rule for histograms (Freedman & Diaconis, 1981). The size-biomass spectrum slopes are mostly positive (>89%) and show strongly different distributions according to the different sampling methods, as they reflect the physical conditions of the habitat itself. Overall the allometric scaling, here as means size-biomass spectrum slope ± Standard Error [SE] with confidence interval as [(5%; 95%) CI], was 0.564 ± 0.354 SE (−0.431; 1.261) CI, and steeper in the larger rivers and shallower in the smaller streams. Water bodies investigated by boat (Method A) had with a mean size-biomass spectrum slope of 0.976 ± 0.191 SE (0.639; 1.301) CI the highest scaling, followed by wading water bodies (Method D) with 0.555 ± 0.287 SE (−0.073; 1.113) CI and finally by longline water bodies (Method E) with 0.377 ± 0.496 SE (−0.751; 1.375) CI as lowest scaling.

Nonparametric analysis is among the best approaches for demonstrating actual differences between scaling groups and true dependencies between our three electrofishing sets. The Mantel test by randomization (Monte Carlo test, 10,000 runs, random number seed 2643) showed a positive association between the allometric scaling matrix and the sampling methods matrix as indicated by all observed Z (sum of cross products) greater than average Z from randomized runs (p = 0.0001, r_M_ = 0.107). Also the calculations performed between the allometric scaling and the sampling matrices using the asymptotic approximation method (Mantel, 1967) show the significant t-value of 13.95 (p < 0.000001). Hence the sampling method itself (here as proxy for the kind of electrofishing chosen according to the water body typology) clearly determines at each location the kind of biological signal detected.

There are not only good methodological reasons to keep these three electrofishing methods separate, but also statistical reasons as shown by the frequency distribution of the slopes. Merging the mass-abundance relationship log(N) = a×log(M) + b into log(B) = log(M) + log(N), we re-obtain the size-biomass spectrum slope log(B) = log(M) + a×log(M) + b = (1+a)×log(M) + b. Hence, these two allometric slopes differ overall by one unit, implying an inverse (negative) mass-abundance relationship in most sites sampled by either method D or E and a direct (positive) mass-abundance relationship in almost all boat-sampled sites (Method A). In particular, at α = 0.05 only 14.1% of the mass-abundance slopes of the boat-sampled water bodies were statistically undistinguishable from the −0.75 coefficient, in contrast to wadeable streams (37.3% of those sampled with Method D and even 52.8% of those sampled with Method E were statistically undistinguishable from either the size-biomass slope of +1/4 or the mass-abundance slope of −3/4). Applying the Pareto rule to the histograms, only 0.88% of all the fish assemblages seem to show steep allometric slopes (Figure 2).

What seems to matter the most is the statistical weight of native predators and introduced omnivores. For instance, large predatory species such as the northern pike (*Esox lucius*) and the muskellunge (*Esox masquinongy*) have been captured strictly by boat electrofishing (as expected seen that large predatory species have a niche in broader and deeper water bodies), whereas the introduced common carp *Cyprinus carpio* has been ubiquitously sampled regardless of the type of electrofishing locally applied. However, the different fresh body-mass average of the *C. carpio* individuals sampled by different types of electrofishing (European carps with Method A share a body-mass average of 2.46 kg, almost three times the average weight of 0.87 kg of the carps with E) is obviously mirroring the variation in habitat type from large rivers to small streams. Wilson (1975) pointed out that a larger competitor can always invade, and a slightly larger competitor would exclude the resident. In the case of the carp, larger aliens in larger rivers, and smaller aliens in smaller rivers, must have equal competitive and invasion success.

Summarizing, much more large-sized fishes are observed in boatable systems than in wadeable streams, where, in turn, much more small-sized individuals are observed. This has clear implications for material flow rates across the higher trophic levels of the aquatic food web. Seen that so often larger individuals consume resources unavailable to smaller competitors, material flow rates across the trophic levels clearly differ between the food webs of boatable systems and those of wadeable streams, as reflected in the size-biomass spectra. Not only the biomass flow, but also the number of fish species change: the biodiversity of wadeable streams in the absence of second-order predatory fishes is in fact much lower than in boatable systems in the presence of second-order predatory fishes. Our results resemble a paradox, as universal laws are per definition regarded as independent from either sampling methods or habitat types. This is clearly not the case here and we show the conceptual menace of integrating data from comparable but still different sites with, to our knowledge, novel computational evidence even within our monophyletic assemblages.

Data processing and data assimilation clearly require new ecoinformatic and analysis tools (Luo et al., 2011), and as Leonelli (2014) wrote, curators have to decontextualize their resources, so that data *“can travel outside of their original context and become available for integration.”* A key principle underlying the design of databases is normalization, although the application of normalization alone is not sufficient. In this paper we urge for a kind of statistical decontextualization before performing data-driven investigations, in our case by showing the extent to which different sampling methods used to reconstruct fish communities strongly and surprisingly influence the allometric scaling law so often assumed to be universal.

**Figure 1.**
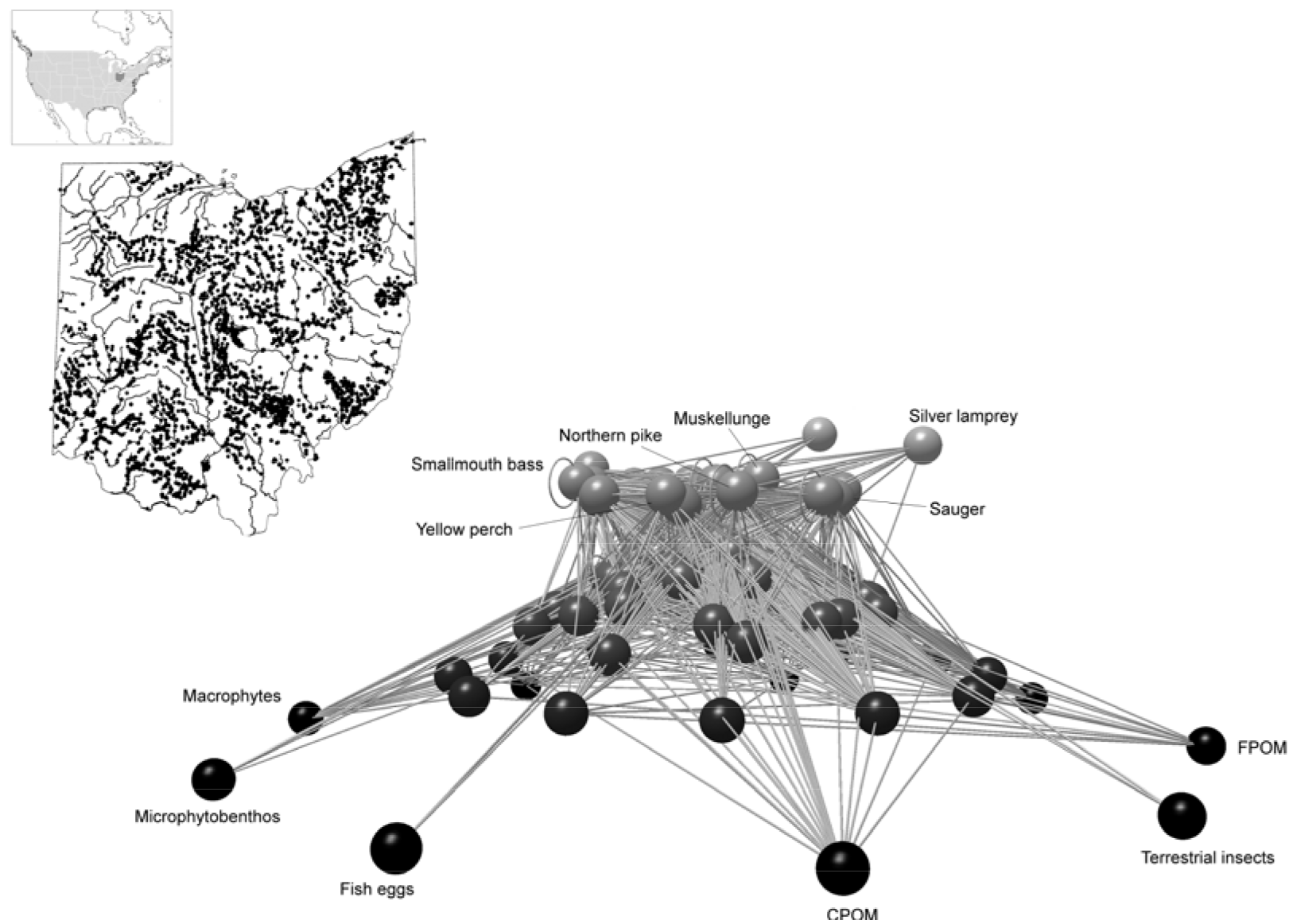
*Network of networks (meta-food web visualized according to Yoon et al., 2004) for all species within 2051 fish communities across the state of Ohio, USA, based on a reduced dataset and the expected trophic links according to the information reported in www.fishbase.org. On average, each species has at regional level 20 possible trophic links. Representative basal resources (e.g., CPOM: coarse particulate organic matter, and FPOM: fine particulate organic matter) and top predators are shown as well.*

**Figure 2.**
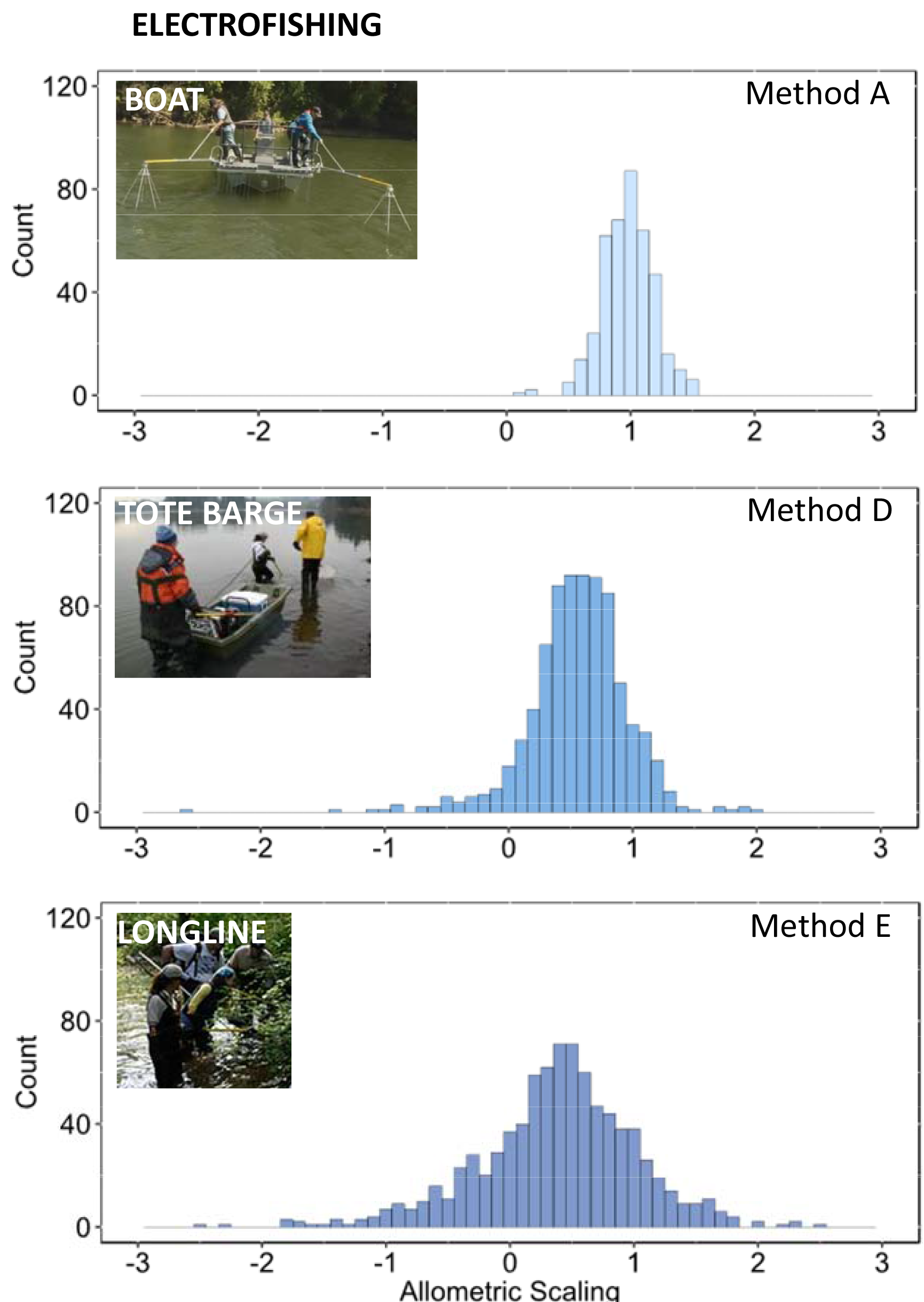
*Shifts of size-biomass spectra across Ohio, USA, according to three independent methodological configurations (electrofishing type).*

## ACKNOWLEDGEMENTS

*We are deeply indebted to Scott Dyer (Procter & Gamble, Cincinnati, Ohio) and Dennis Mishne (US-EPA Ohio) for supporting our data mining and to Henri Den Hollander and Valentina Sechi (RIVM) for their feedback. This research was partly supported by the EU (SOLUTIONS grant no. 603437 to CM)*.

